# Host niche breadth differentially modulates the effects of anthropogenic disturbance across generalist and specialist parasites

**DOI:** 10.1101/2025.05.11.653240

**Authors:** Ashwin Warudkar, R Gayathri, Chiti Arvind, Farah Ishtiaq, Guha Dharmarajan, V.V. Robin

## Abstract

Anthropogenic disturbances in natural habitats increase the risk of emerging infectious diseases in free-ranging host communities. Hence, understanding the processes driving such patterns is critical towards One Health. While anthropogenic disturbance is known to promote habitat-generalist host species through biotic homogenisation, generalist parasites (wider host breadth) also respond positively to the disturbed habitats. We hypothesise that generalist parasites are more likely to infect generalist hosts. We tested this hypothesis in a sky island system where generalist (*Plasmodium*) and specialist (*Haemoproteus*) haemosporidian parasites infect a range of bird hosts in disturbed and natural forest patches.

We used a natural experiment framework to control for climatic differences (similar elevation) and habitat quality (same habitat type, but varying disturbance matrix). We collected 1106 samples from the field and examined the genus-level parasite prevalence and the host specificity of individual parasite lineages. Our results suggest that the generalist (*Plasmodium*) parasites are more prevalent in generalist birds, irrespective of disturbance. However, the specialist (*Haemoproteus*) parasites were more prevalent in specialist birds in natural forests than in disturbed forests. Among the individual parasite lineages, we found the host specificity to be associated with the degree of habitat specialisation of their host species. Our results provide evidence for the tendency of generalist parasites to infect generalist host species – a potential mechanism for a higher risk of emerging infectious diseases in human-dominated regions. We emphasise the role of host ecology in understanding the impact of anthropogenic disturbance on parasite prevalence in free-ranging host communities.

## Introduction

Globally, the implications of landscape and climate change are numerous; additionally, emerging infectious diseases are creating unprecedented challenges for biodiversity conservation. The pathogen outbreaks can cause drastic, stochastic population changes within a short time (Daszak et al. 2000, De Castro and Bolker 2005, Cunningham et al. 2017, Grimaudo et al. 2024). Additionally, the knowledge gap regarding parasites’ specificity makes the host response unpredictable during an outbreak (LaDeau et al. 2011, Altizer et al. 2013). Environmental factors, such as habitat and climate change, affect the free-ranging host species (Travis 2003, Jeltsch et al. 2011, Mantyka-pringle et al. 2012, Frishkoff et al. 2016). The host-dependent parasites in such scenarios may show changes in their prevalence in their hosts (Cable et al. 2017, Gsell et al. 2023). Many parasites show high specificity (specialist) to one or a group of closely related host species (Shaw et al. 2020). The exceptions are those with higher host breadth (generalist), comprising phylogenetically distant host species (Shaw et al. 2020). The host species, similarly, can have narrow or broad niche breadths with respect to their habitat choice and spatial distribution (Carscadden et al. 2020). Their degree of habitat specialisation would drive their differential response to environmental change, with the specialist species being more vulnerable (Dennis et al. 2011, Ducatez et al. 2014). Environmental changes may thus impact the parasites indirectly, depending on their interactions with free-ranging hosts. Understanding these impacts, e.g. changes in parasite prevalences in a host species community and the specificity of their interactions, becomes crucial for risk assessment of potential outbreaks.

Anthropogenic disturbance is known to cause biotic homogenisation of free-ranging vertebrates (McKinney 2006, Sidemo-Holm et al. 2022), wherein the habitat generalists can survive with disturbance, promoting their relative abundance (Vargas Soto et al. 2022, Yang and Hu 2025). Vertebrate reservoir hosts of zoonotic parasites highlight this pattern (Gibb et al. 2020). On the other hand, the human disturbance is also shown to favour generalist parasites (Budria and Candolin 2014, Dharmarajan et al. 2021, Prati et al. 2022, Wilson et al. 2024). To our knowledge, no study has described the mechanism connecting generalist hosts and generalist parasites. For instance, we suspect that the generalist parasites will likely infect generalist hosts. This may allow them to escape a potential dilution effect through host switching and thrive in the altered habitat. Hence, the degree of hosts’ habitat specialisation should explain the spatial variation in the prevalence of generalist parasites.

Avian haemosporidia are blood parasites of birds transmitted by dipteran vectors and are widely studied host-parasite systems. The most commonly found genera include *Plasmodium*, *Haemoproteus* and *Leucocytozoon* (Lapointe et al. 2012, Valkiūnas and Iezhova 2017, Fecchio et al. 2020). Avian haemosporidian parasites came to the limelight after the introduction of *Plasmodium*, and its subsequent outbreak in the Hawaiian endemic birds caused local extinctions (Warner 1968, van Riper et al. 1986, Lapointe et al. 2012, Samuel et al. 2015). *Plasmodium* is broadly considered a generalist in terms of its host breadth (Doussang et al. 2021). In contrast, *Haemoproteus* is considered a specialist, infecting fewer or closely related host species, although variations exist within members of both genera (Valkiunas 2004). The specificity and coevolution of these parasites with their hosts are essential yet understudied factors in determining variation in the parasite infections across an environmental gradient.

Host diversity can have a positive (amplification) or a negative (dilution) effect on parasite prevalence (Halliday and Rohr 2019, Ferraguti et al. 2021). However, in the case of vector-borne infections, the vector diversity too can exhibit either effect on parasite prevalence (Randolph and Dobson 2012, Kocher et al. 2023). Habitat fragmentation is one of the common ways human disturbance is characterised at a landscape scale, wherein bird diversity is generally expected to decrease in disturbed habitats (Srinivasan 2013, Sam et al. 2014, Bełcik et al. 2020). Hence, the haemosporidian prevalence will decrease with disturbance if it experiences an amplification effect or will increase if it experiences a dilution effect. Nonetheless, variations exist in how different parasite genera and lineages are impacted by environmental change (Ferraguti et al. 2021), a pattern not well studied across different ecosystems.

Although oceanic island birds have been shown to experience high risk from avian haemosporidia, the continental sky islands or the montane habitat islands remain underexplored (Samuel et al. 2011, Lapointe et al. 2012, Niebuhr et al. 2016) while also being considered vulnerable to climate change and facing rapid landscape change (Arasumani et al. 2019, Love et al. 2023). We studied the haemosporidian parasite community in Palani-Anamalai hills (10 24” N - 10 01” N and 76 58” E - 77 40” E), one of the Shola Sky Islands of the Western Ghats of India, typically above 1400 meters above (mean) sea level (m.a.s.l.). Like many other sky islands, the local montane *shola* bird community consists of a few endemic, elevationally restricted species (Robin & N 2012, Love 2023) and many species with distributions across a broader elevational gradient. This landscape has changed rapidly (up to 60%) with woody invasives and human settlements in the past century (Arasumani et al. 2018, 2019, 2021). The shola forests in the modified landscape exist as isolated patches with various forms of human impacts and are surrounded by anthropogenic habitats, including human settlements. Previous studies on avian haemosporidian parasites (*Plasmodium* and *Haemoproteus*) in the Shola Sky Islands described broad biogeographic patterns of their diversity and prevalence across mountains (Gupta et al. 2019). However, explicit tests of human disturbance connecting parasites’ specificity across a gradient of hosts’ habitat specialisation have not been conducted. The response of both *Plasmodium* and *Haemoproteus* to disturbance or undisturbed forests is known to be variable across different geographies (Bonneaud et al. 2009, Chasar et al. 2009, Loiseau et al. 2010, Laurance et al. 2013). Hence, testing this effect with intensive sampling accounting for host ecology may reveal critical processes and add to the global understanding of parasite ecology. This study assesses the relationship between generalist hosts and generalist parasites with anthropogenic disturbance. We explicitly examine the host specificity of individual parasite lineages across the gradient of habitat specialisation of hosts to decipher the relationship between generalist hosts and generalist parasites. We expect parasites that infect phylogenetically diverse host species (generalist parasites) to be more common in host species that thrive in diverse habitats (habitat generalist hosts).

## Methods

### Field Sampling & disturbance

We sampled the bird community in the sky island landscape (>1400masl) of the Palani-Anamalai Hills, Western Ghats, India (Figure 1A). To control for the effect of land cover, we conducted our sampling only inside natural shola forests and between winter and summer (November to March), considering the stable and aseasonal climate in the tropical sky islands of the southern western ghats (Bose et al. 2016, Page and Shanker 2020). The breeding of birds in this landscape may occur throughout the year, with a peak from December to June (Ramesh et al. 2012, Jha 2023). We conducted field sampling at six sites between 2020 to 2023. We followed a natural experimental set-up where we sampled the same habitat type – shola forests, but these occurred under two landscape scenarios: 1) those close to human-modified landscapes (e.g. large settlements within 1 km) considered as disturbed forests: Kodaikanal (2100masl), Munnar (1550masl)and Devikulam (1675m); and 2) those situated within protected areas and far from human-modified landscapes (> 5km) considered as natural forests: Kookal Shola-Anamalai Tiger Reserve (1925masl), Berijam-Kodaikanal Wildlife Sanctuary (2250masl) and Bandar - Shola National Park (2300masl) (Figure 1A). Thus, we were sampling a shola forest patch either surrounded by human habitation as disturbance treatment or nested within an extensive shola forest as the natural control. Such a design was used to control for a variation in the community of hosts and roughly similar habitat structure. We used mist nets to sample the local bird community, collected blood samples from the brachial vein, and stored them in Queen’s Lysis Buffer or on FTA cards (Qiagen Whatmann and HiMedia). We also measured standard morphometric measurements of birds, including tarsi, wings, beak, tail, weight and age (adult vs 1st year juvenile) of the bird. We used 10 to 30 mist nets at each site, sampled for 10-15 days in one session, and sampled across three years. Some sites (Kodaikanal and Bandar) were sampled multiple times to achieve larger sample sizes within a year. Our study species comprised 58 species, most of them common to the understory bird community in the Shola Sky Islands.

**Figure 1.**
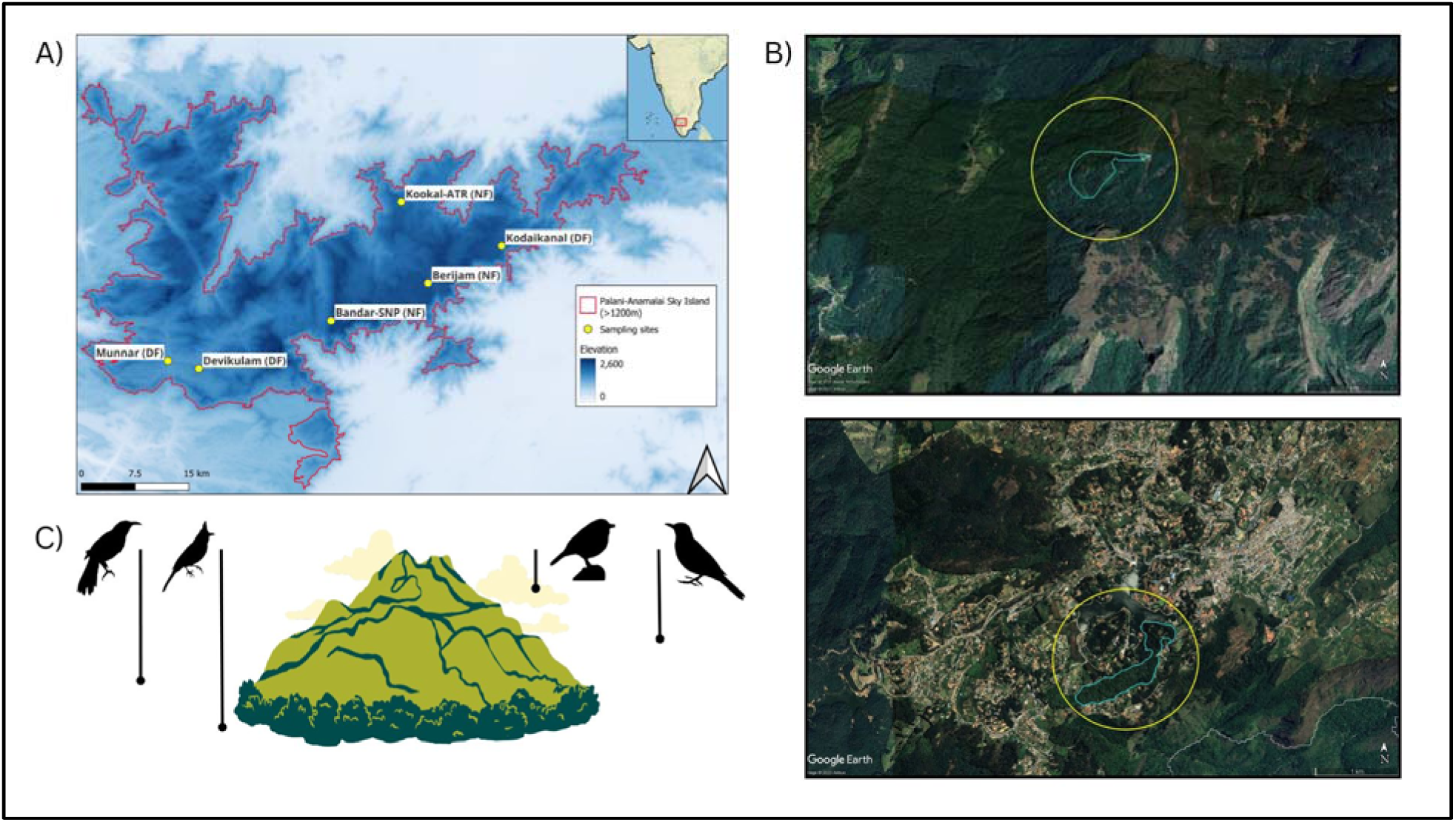
A) Study area with sampling locations (DF: disturbed forest; NF: natural forest). B) Satellite imagery of Bandar-SNP, a natural forest site (above) and Kodaikanal, a disturbed forest site (below); blue outline inside the circle indicates the sampled forest patch; C) Illustration of the lower elevation limit of the bird species on a mountain as a measure of habitat specialisation.

### Parasite detection and identification

DNA was extracted (Qiagen DNAesy Blood and Tissue Kit), and two different PCR techniques were used to detect the presence of haemosporidian parasites. We simultaneously amplified genus-specific regions for three parasites, *Plasmodium*, *Haemoproteus* and *Leucocytozoon*, using a Multiplex PCR protocol (Ciloglu et al. 2019) and detected the parasite infection through Agarose Gel Electrophoresis. We then performed nested PCR (Waldenström et al. 2004, Hellgren et al. 2004) and amplified mitochondrial Cytochrome B gene fragments from every positive sample. We sequenced (Sanger) the PCR-positive amplified product and assembled forward and reverse reads (Geneious Prime 2024.0). The parasite genus and lineage were identified using GenBank using the Nucleotide BLAST algorithm (https://blast.ncbi.nlm.nih.gov/Blast.cgi) and MalAvi database BLAST tool (Bensch et al. 2009). Unidentified sequences with one or more base pair differences (different haplotypes) were treated as new lineages.

### Host-parasite co-phylogeny to determine genus-level parasite specialisation

We aligned all the DNA sequences using the Muscle alignment (Edgar 2004). The best-fitting partition scheme and the best substitution model were determined using PartitionFinder (Lanfear et al. 2012). Bayesian phylogenetic analyses were performed in Mr Bayes (Ronquist and Huelsenbeck 2003). Two independent runs, each with four MCMC (Markov Chain Monte Carlo) chains, were performed for 20 million generations and sampling every 500^th^ generation by discarding 25% of samples as burn-in. To evaluate the cophylogenetic relationship between hosts and parasites, we used ParaFit (Legendre et al. 2002) to evaluate the congruence between host and parasite phylogeny based on an association matrix. A global test statistic (Legendre et al. 2002) was computed to measure the overall congruence between the phylogenies.

### Hosts’ environmental specialisation and Parasites’ host breadth

Specialist species in sky islands occur on mountaintops (Robin and Nandini 2012, Robin et al. 2015). Elevation becomes a surrogate for various environmental factors limiting the occurrence of bird species ((Robin and Nandini 2012, Robin et al. 2015). For all our study species, we calculated the lower elevation limit in the southern Western Ghats using citizen science eBird (eBird 2021) records, filtered for the effort less than 1km, to minimise elevational inaccuracy bias. The log-scaled values of this limit were considered a metric of the host’s environmental specialisation, where higher values (species restricted to the mountain tops) indicated a more elevationally restricted/specialised bird species. Elevational specialisation is a widely recognised ecological niche specialisation in many mountain systems and can result from various environmental and physiological or ecological (competition; predation) processes (Chen et al. 2024). For our study species, too, elevational breadth or lower elevation limit of occurrence represented habitat specialisation (calculated from the species’ usage of different habitat types and derived from (Bellwood et al. 2006)’s approach; see supplementary material and Fig S1). We calculated each parasite lineage’s phylogenetic host breadth (mean phylogenetic distance between the infected host species) as a metric of parasite specialisation. The global host breadth was considered to account for the known fundamental host breadth of the parasite as opposed to the local host breadth. We accessed the MalAvi dataset (MalAvi database, (Bensch et al. 2009), accessed on 5 November 2024) to obtain the global host diversity for each parasite lineage in the present study. The host phylogeny was created by subsetting the global bird phylogeny (McTavish et al. 2024). The mean phylogenetic distance between all the hosts globally of a parasite was calculated and considered as parasite specialisation.

### Parasite prevalence among sites

We pooled the parasite detections from both multiplex PCR (Ciloglu et al. 2019) and nested PCR (Hellgren et al. 2004) methods and used the consolidated detections as presence/absence for each sample which allowed us to account for false negatives. We used binomial mixed effect regression models (GLMM) using ‘lme4’ package in R to model *Plasmodium* and *Haemoproteus* prevalence. The parasite prevalence was modelled against human disturbance (categorical: disturbed forests vs natural forests) and the hosts’ environmental specialisation (continuous: log of lower elevation limit), along with their interaction term in a complete model. The host species identity, host family, and age of the individual were considered random effects (random intercept and slope). We also estimated the body condition of every individual bird as a scaled mass index proposed by ((Peig and Green 2009)) and tested for its effect on the prevalence of parasites. The models with all combinations of independent variables were compared, and the best model was selected with the lowest AICc score.

### Host-specificity of Parasite lineages vs Host’s environmental specialisation

The global phylogenetic host breadth of parasites was correlated with the mean of log-scaled lower elevation limit of the birds they infected. This allowed us to understand the infection tendency of parasites in bird species with different degrees of specialisation. Parasite lineages infecting only one host species were omitted.

## Results

### Haemosporidian prevalence among hosts and sites

We sampled 1106 individual birds (596 within natural forests and 510 birds within disturbed forests) belonging to 58 species from six localities across the study area. 12 species were winter migrants. However, these migrant birds also spend nearly 8 months in this landscape, and hence, their samples were included in the analysis. We found 517 birds (505 adults and 12 juveniles) from 36 species infected with at least one of the three haemosporidian parasite genera (prevalence 50.54%) across the landscape. *Haemoproteus spp.* were the most prevalent among all parasite genera, infecting 395 birds from 27 species (prevalence 35.71%), followed by *Plasmodium spp.* Infecting 134 birds from 31 species (prevalence 12.11%), and *Leucocytozoon* infecting only 33 birds from 15 species(prevalence 2.98%). The prevalence of haemosporidian parasites varied across bird species. *Plasmodium* was found most dominantly in the Indian blackbird, *Turdus simillimus* (prevalence: 87.09%; N=62), followed by the Indian scimitar babbler, *Pomatorhinus horsefieldii* (prevalence: 33.33%; N=39). *Haemoproteus* was found most dominantly in Indian White-eye, *Zosterops palpebrosus* (prevalence: 85.5%; N=200), followed by the Palani laughingthrush, *Montecincla fairbanki* (prevalence: 58.9%; N=151).

### Effect of human disturbance on the haemosporidian prevalence

The detection accuracy of parasite infection differed between multiplex PCR (Ciloglu et al. 2019) and nested PCR (Hellgren et al. 2004). Of the total infected birds, 92.45% of infections were detected through multiplex PCR, 88.58% of the infections were detected through nested PCR, and 81.04% were detected commonly by both methods (McNemar’s Chi-squared test Chi-squared=4.08; p-value=0.04) (see supplementary Table S1 for details). The prevalence of *Leucocytozoon* parasites was very low, and since our interests were in the communities of parasites and hosts, we excluded them from further analysis. We found an overall cumulatively higher prevalence of all the haemosporidian parasites in the natural forest sites (50.58%) than in the disturbed forest sites (41.96%) (Chi-squared=8.34; p-value=0.002) the difference was driven by prevalence of *Haemoproteus*. Through separate binomial GLMM fit to the *Haemoproteus* and *Plasmodium* prevalence, we found different variables influencing the prevalence of generalist *Plasmodium* and specialist *Haemoproteus* parasites (Figure 2). The best model explaining *Plasmodium* prevalence indicated that the environmental specificity (lower elevation limit) of the host species had a negative effect on *Plasmodium* prevalence (Odds ratio: 0.77; ±SE=-0.26±0.12; p-value=0.029) (Figure 2A). However, there was no statistical difference in *Plasmodium* prevalence between the disturbed and natural forests (Odds ratio: 1.48; (disturbed)±SE=0.38±0.25; p-value=0.119). In the case of *Haemoproteus*, an interactive effect of the site variable, viz., disturbance and the host’s environmental specificity (lower elevation limit), best explained its prevalence pattern. The lower elevation limit of the host species had a negative effect on the *Haemoproteus* prevalence in the disturbed forests (Odds ratio: 0.62; (interaction)±SE=-0.48±0.08; p-value=2.27e^-08^) (Figure 2 B). This interactive relation was typically seen in the birds endemic to high elevations, e.g. The White-bellied Sholakili, *Sholicola albiventris,* and Palani Laughingthrush; both had a considerably higher prevalence in the natural forest sites than the disturbed forest sites (see Figure 3 A & B). The *S. albiventris* population in disturbed forests are nearly free of infection, while those in natural forests (except Berijam) have 44.3% and 38.5% prevalence. The individual bird’s body condition and age did not explain any variation in the prevalence between disturbed and natural sites for either parasite genera. Furthermore, generalist birds (as described by lower elevation limits–see methods for details) are captured more in disturbed forests than in natural forests (Mann-Whitney U test: p-value=0.001).

**Figure 2:**
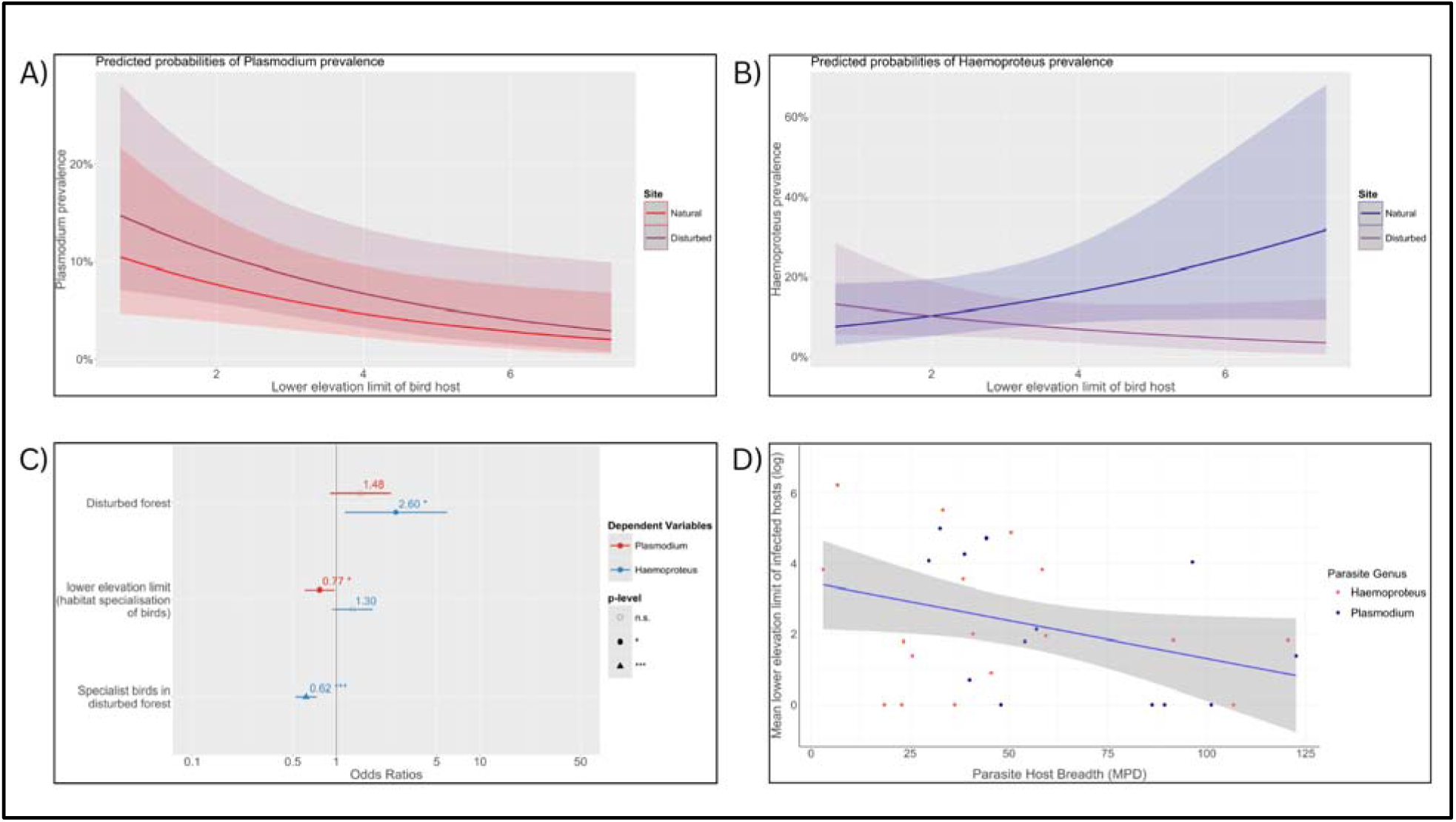
A & B) Effect of host specialisation (lower elevation limit) on the prevalence (predicted probabilities) of *Plasmodium* and *Haemoproteus* parasites between natural forests and disturbed forests. C) Coefficients (odds ratio) of binomial mixed effect models for the effect of disturbance and host specialisation on *Plasmodium* and *Haemoproteus* prevalence. D) Relationship between host specialisation (phylogenetic host breadth) of parasite lineages and th habitat specialisation of the host they infect, depicting the tendency of host-generalist parasites to infect habitat-generalist hosts.

**Figure 3:**
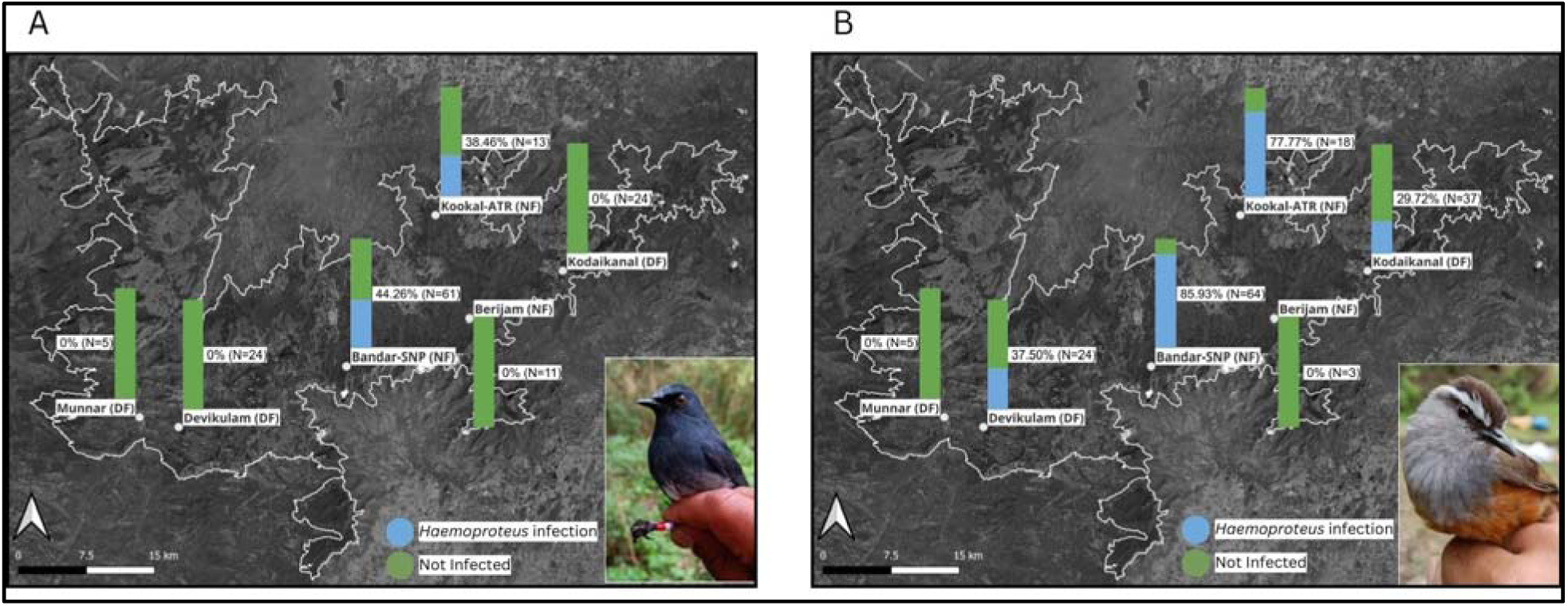
Prevalence of *Haemoproteus* in the habitat specialist hosts *Sholicola albiventris* (A) and *Montecincla fairbanki* (B) across disturbed forests (DF) and natural forests (NF).

### Haemosporidian lineage diversity and host specificity

We sequenced 479bp mitochondrial Cytochrome B gene fragments from 428 birds of 30 species. We identified 53 unique lineages of haemosporidian parasites, of which 21 lineages belonged to *Plasmodium spp.* and 32 belonged to *Haemoproteus spp.*, identified across 17 and 25 bird species. We discovered 11 new lineages and newly recorded 14 lineages from the Western Ghats, updating the known regional richness to 72. The disturbed forest sites had more haemosporidian lineages (38; 16 *Plasmodium* and 32 Haemoproteus) than the natural forests (32; 12 *Plasmodium* and 20 *Haemoproteus*). Disturbed and natural forests shared 17 lineages and exclusively had 21 and 15 lineages. However, there was no difference in the mean phylogenetic host breadth of both parasite genera between disturbed and natural shola forests.

We compared the hosts’ mean phylogenetic distance (MPD) between two parasite genera at local and global scales and found contrasting results. We found a negative correlation between the mean phylogenetic host breadth of parasite lineages and the mean lower elevation limit (habitat specialisation) of the hosts they infect (Figure 2D). The parasite community in disturbed forests had a higher average phylogenetic host breadth than that in natural forests (Mann-Whitney U Test; p-value<0.01). *Plasmodium* had a higher global mean phylogenetic host breadth than *Haemoproteus*, but the difference was not significant for the local phylogenetic host breadth (t-test ; p value>0.05).

Together, members of Muscicapidae (e.g. Nilgiri Flycatcher *Eumyias albicaudatus*, Black and Orange Flycatcher *Ficedula nigrorufa*), followed by Turdidae (e.g. *Turdus simillimus*, Nilgiri Thrush *Zoothera neilgherriensis*), had the highest parasite lineage richness within the study area. The two most range-restricted endemic hosts differed in their parasite lineage richness. However, most of the *Haemoproteus* lineages were singletons (infected only one host species), the lineage ‘COLL2’ infected 7 species (Figure S3) in the study area and is also a geographically widespread lineage. We found 21 new host associations for 11 parasite lineages, updating host breadth and geographic distribution of many singletons, particularly for the *Haemoproteus* lineages MONMER01 and SHOMAJ03, which were previously known only from the Ashambu hills and the Nilgiri hills.

## Discussion

In this study, we intensively sampled the cloud forests in a sky island landscape to understand the impact of human disturbance on the haemosporidian prevalence in birds and the underlying mechanism driving the prevalence patterns. We show that the host’s environmental specialisation plays a role in determining the response of generalist and specialist parasites to human disturbance. *Plasmodium* prevalence depended on the host’s habitat specialisation, which was more prevalent in the generalist host species. In contrast, *Haemoproteus* prevalence differed across hosts’ specialisation between disturbed and natural forests, wherein specialist hosts are more likely to be infected in natural habitats than disturbed ones. Further, the generalist parasite lineages (with higher phylogenetic host breadth) infected the generalist host species (with broader elevational niches).

Our results highlight how hosts’ habitat specialisation interacts with human disturbance to determine the prevalence of parasites in mountaintop ecosystems. Our study of shola forest species adds to the literature of habitat specialisation based on the elevational breadth of a species (Peh et al. 2012, Mermillon et al. 2021, Korejs et al. 2025). We demonstrate that using the lower elevation limit (elevational breadth) as a proxy for hosts’ habitat specialisation explains the variation in parasite infection rates amidst the anthropogenic changes. The prevalence and diversity of avian haemosporidian parasites have been assessed for different host traits, including sociality/flocking, sexual dimorphism, incubation period, body size and participation in mixed flock (Arriero and Møller 2008, Santiago-Alarcon et al. 2016, Rodríguez-Hernández et al. 2021, Fecchio et al. 2021a) as well as individual bird traits such as body mass, body condition, fluctuating asymmetry (Santiago-Alarcon et al. 2016, van Hoesel et al. 2020, Gupta et al. 2020). However, studies on host communities that consider the host’s habitat or elevational specialisation are limited (Ferreira Junior et al. 2017). Our study also gained from an explicit study design where our selection of sites controlled for the habitat type –forests, and examined disturbance as a function of the anthropogenic matrix surrounding the sites (forest patch).

*Plasmodium* parasites are usually found in a wide range of hosts and are considered generalists (Doussang et al. 2021). In the Shola Sky Islands, we show that habitat generalist hosts, occurring across a wider elevational range, are more likely to be infected by *Plasmodium*, such as in (Loiseau et al. 2012, Clark et al. 2015), but not anthropogenic disturbance, unlike (Hernández-Lara et al. 2017, Fecchio et al. 2021b, Dharmarajan et al. 2021). This mechanism could underlie the indirect effect of human disturbance on the *Plasmodium* infection rate, although not observable directly at intermediate disturbances. *Haemoproteus,* on the other hand, generally exhibits a greater degree of host specificity and higher lineage diversity and, in turn, is broadly considered a specialist genus (Doussang et al. 2021). We show that generalist birds were more likely to be infected by *Haemoproteus* in disturbed forests; in contrast, specialist birds were more likely to be infected in natural forests. This effect is prominently reflected in the prevalence of three lineages in two specialist birds (COLL2 in *S. albiventris* and MONFAI02 and MONMER01 in *M. fairbanki*). The higher prevalence of *Haemoproteus* in natural or undisturbed forests is reported in tropical Australia (Chasar et al. 2009, Laurance et al. 2013) and African rainforests (Chasar et al. 2009, Laurance et al. 2013). *Haemoproteus* prevalence has been known to correlate with forest cover (Fecchio et al. 2021b), likely following the distribution patterns related to its vectors, *Culicoides* flies, which are also associated with natural forests in the Neotropics (Loaiza et al. 2019, Mendez-Andrade et al. 2024). Assessing *Culicoides* vector diversity across disturbed and natural forests in the Western Ghats sky islands is the key to explaining our observed pattern.

Despite controlling for habitat type in our study design, our host sampling indicated a proportionally higher abundance (and sampling) of generalist hosts in disturbed forests than in natural forests (Figure S2). In this circumstance, greater *Plasmodium* prevalence in generalist hosts potentially supports the dilution effect (Halliday and Rohr 2019), where the loss of diversity due to disturbance increases generalist species’ relative abundance (Mayfield et al. 2010, Clavel et al. 2011, Matuoka et al. 2020); (Freedman et al. 2010, Clavel et al. 2011). On the contrary, the higher *Haemoproteus* prevalence in specialist birds in natural forests indicates an amplification effect.

In our study, the scale of examination of the parasite specificity indicated the outcome of the relationship of association with the hosts. The global, but not local (within the study area), host breadth of parasite lineages showed a negative relationship with the host’s environmental (elevational) specialisation. Our study indicates that generalist parasites infect generalist hosts when global host breadths of parasites are considered, such as in (Loiseau et al. 2012), unlike the local host breadth considered by (Huang et al. 2018). The global host breadth captures the fundamental infection potential of a parasite lineage. Several generalist (higher host breadth) parasite lineages in our study have diverse hosts across their global distribution range, yet many such lineages may infect only a few host species locally (Huang et al. 2018). Nonetheless, their potential for host switching can play a role in the face of contemporary environmental changes. The parasites sampled in disturbed forests had slightly higher phylogenetic host breadth than those in natural forests. The future increase in habitat disturbance may further facilitate the spread of generalist parasites in this montane bird community.

Considering the broader significance of our study, we demonstrate that the process of generalist parasites likely infecting generalist hosts can be a mechanism behind anthropogenic disturbance driving the prevalence of generalist parasites. The generalist parasites have a higher host-switching potential reflected in their wider host breadth, while specialist parasites invest in a particular host or closely related group of hosts (Agosta et al. 2010). The anthropogenic disturbance can cause the decline of specific hosts of specialist parasites (Cizauskas et al. 2017, Cable et al. 2017). In contrast, generalist parasites are more adaptable and can exploit the opportunity provided by habitat generalist hosts. We document this process to occur both within the parasite genus as well as the individual parasite lineages in the haemosporidia of birds.

As with many other studies of avian malaria, one of the challenges that remains with our study is the undocumented role of vectors. The apparent low prevalence of *Haemoproteus* in habitat specialist birds from disturbed forests may be mediated by vectors, which were out of the scope of the present study. The *Culicoides* flies are shown to respond differently to human disturbance (Loaiza et al. 2019, Mendez-Andrade et al. 2024) and may show specificity to parasites or vertebrate hosts (Martínez-de la Puente et al. 2015, Bukauskaitė et al. 2019). On the other hand, specialist parasites like *Haemoproteus* may exhibit a density-dependent transmission at lower host densities, like in human malaria (Villena et al. 2024), resulting in their low prevalence in specialist birds with disturbance. The avian haemosporidia are complex host-parasite-vector systems (Valkiunas 2004). Moreover, habitat modification could change microclimatic parameters affecting parasite phenology in vectors and hosts (Mozaffer et al. 2022). A longitudinal study involving vector sampling at the same spatial scale would help tease apart the temporal variation in prevalence patterns.

Mountain systems are among the most vulnerable systems undergoing rapid anthropogenic change, with mountaintop-restricted endemic species at risk of extinction (Schmeller et al. 2022). Understanding host-parasite associations and the infection tendency of generalist and specialist parasites is crucial in assessing disease outbreak risk amid global and local change. In this study, we demonstrate that human disturbance in isolation may not explain the difference in the infection probability across generalist and specialist parasites unless it is assessed through the ecological lens of the host’s environmental specialisation. Especially with host-specific parasites, human disturbance in interaction with host ecology may provide a deeper understanding. We also argue that the scale of the system matters when such patterns and mechanisms, e.g. dilution vs amplification, are tested (Keesing et al. 2006, Moens and Pérez-Tris 2016, Ferraguti et al. 2021). Taken together, our study shows axes of diversity in which hosts and parasites can interact in a sky island landscape.

## Acknowledgement

We thank the state forest departments of Tamil Nadu (Kodaikanal forest division: S.N. Thejaswi, P.K. Dileep, Yogesh Kumar Meena, Anamalai Tiger Reserve: I. Anwardeen) and Kerala (Eravikulam National Park: S.V. Vinod, Munnar forest division: Ramesh Bishnoi) for providing necessary permission to conduct fieldwork. We also thank the range forest officers, section forest officers, beat officers and watchers from all the study sites for their help with field logistics. We thank our field assistants Subhash, Kamraj, Kovai, Yuvraj and Karnan for their help during fieldwork. We are grateful to Vinay K.L., Suyash Sawant, Viral Joshi, Akshay Herur, Arpitha G.P., Aparna Rao, Shivam Shinde, Mohammed Mubin, Yash Deshpande and Karthik Sharma for assisting in the field work and lab work. We also thank Naman Goyal, Vinay K.L., Jobin Varughese, Archita Sharma, Meghana Natesh, Arunima Jain, Akshay Herur, and Madhavi Shukla for valuable discussion and help during the analysis. We thank Kodaikanal International School for field station and logistics support. We thank Uma Ramakrishnan, Ansil B.R., Trevor Price and Ravinder Sehgal for their suggestions on study design, analysis and conceptualisation. This study was funded by the Ministry of Environment, Forest and Climate Change, Government of India, and the National Geographic Society.

## Funding

This study was primarily funded by the Ministry of Environment, Forest and Climate Change (MoEFCC), Government of India’s research grant (reference no. 19-22/2018/RE). This study was also supported by the National Geographic Society Level-2 grant (NGS-93271R-22) awarded to VVR, IISER Tirupati internal funding and Krea University internal funding.

## Authors’ contribution

Conceptualisation: AW(lead), FI, GD(lead), VVR(lead)

Data curation: AW(lead), GR

Formal analysis: AW(lead), GR

Funding acquisition: VVR

Investigation: AW(lead), CA*, VVR*

Methodology: AW*, FI*, GD*, VVR*

Project administration: VVR Resources: VVR

Supervision: VVR(lead), FI(lead), GD(lead)

Visualisation: AW(lead), GR

Writing - original draft: AW(lead), VVR

Writing - review & editing: AW(lead), FI, GD(lead), VVR(lead)

## Competing interests

We declare no competing interests.

## Notes

### Competing Interest Statement

The authors have declared no competing interest.

