## Supplementary Information for "Host niche breadth differentially modulates the effects of anthropogenic disturbance across generalist and specialist parasites"

Parasite detection accuracy of two PCR methods

We found discrepancies in the detection accuracy of parasite infection by either of the PCR methods we implemented (see methods). Out of all the 1107 samples tested, multiplex PCR (Ciloglu et al. 2019) detected 129 *Plasmodium*(11.66%), 365 *Haemoproteus* (33.00%) and 17(1.53%) *Leucocytozoon* infections, while the nested PCR (Hellgren et al. 2004) found 108 *Plasmodium* (9.76%), 368 *Haemoproteus* (33.27%), and 22 *Leucocytozoon* (1.98%) infections. Among 129 *Plasmodium* infections detected by multiplex PCR, nested PCR failed to detect 26 samples (20.15%). In comparison, out of 108 *Plasmodium* infections detected by the nested PCR, the multiplex PCR failed to detect five samples (4.62%). Both nested and multiplex PCRs detected 103 *Plasmodium* infections. For *Haemoproteus*, among 365 infections detected by the multiplex PCR, nested PCR failed to detect 27 samples (7.39%), while among 368 infections detected by nested PCR, multiplex PCR failed to detect 30 samples (8.15%). Both the PCR methods detected 338 *Haemoproteus* infections. For *Leucocytozoon*, the nested PCR failed to detect 11 infections out of 17 (64.70%) detected by the multiplex PCR. However, multiplex PCR failed to detect 16 infections out of 22 (72.72%) by the nested PCR, while both protocols detected six infections (see Table 1).

Table S1. Comparative accuracy of Multiplex PCR and Nested PCR in detecting the haemosporidian infection

| Parasite Genus | Multiplex PCR | Nested PCR | both | Total |
|---|---|---|---|---|
| *Plasmodium* | 129 | 108 | 103 | 134 |
| *Haemoproteus* | 365 | 368 | 338 | 395 |

| | | | | |
|---|---|---|---|---|
| *Leucocytozoon* | 17 | 22 | 6 | 33 |
| Total | 478 | 458 | 421 | 517 |

Using host occurrences across habitats to measure their specialisation

We downloaded eBird occurrence data for all the host species across elevation and Palani-Anamalai hills southwards. We filtered it for the effort less than 1km, to minimise elevational inaccuracy bias. We extracted the data on the vegetation class from (Roy et al. 2015) for all the occurrence points of a bird species. The shola forest and shola grassland habitats from (Roy et al. 2015) are interspersed at a finer scale, and their accuracy is low (personal observation), hence we pooled the bird occurrences across both classes and merged them into a single class 'shola habitat'. We also attached each species' lower elevation limit and elevational breadth (maximum elevation minus lower elevation) to the occurrence dataset. We then estimated the species habitat specialisation derived from (Bellwood et al. 2006) by plotting a Principal Co-ordinate Analysis (PCoA) plot of the normalised variables of species' habitat occurrences and elevational limits and taking the distance of each species from the centroid. The closer the species to the centroid, the more generalist it is and vice versa. We then correlated this habitat specialisation index with the lower elevation limit to validate its use in the study (see Supplementary figure S1)

Supplementary figures

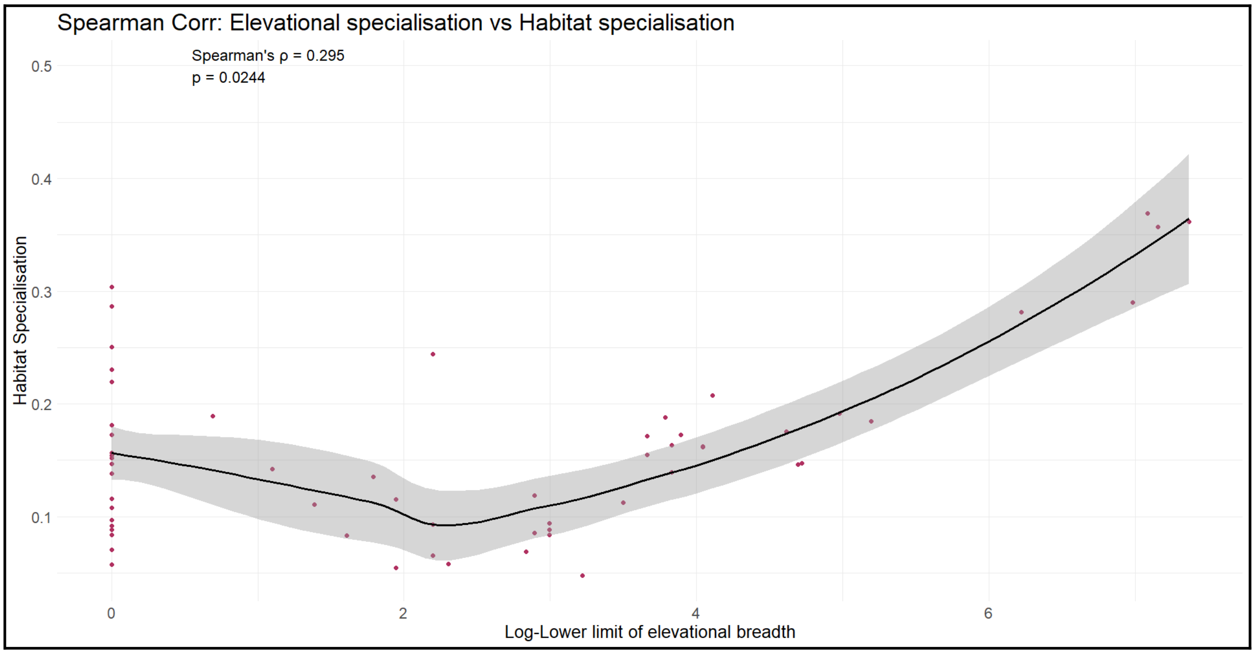


Figure S1.: The Spearman correlation plot of the Habitats specialisation vs log of lower elevation limit of the bird species sampled in the present study.

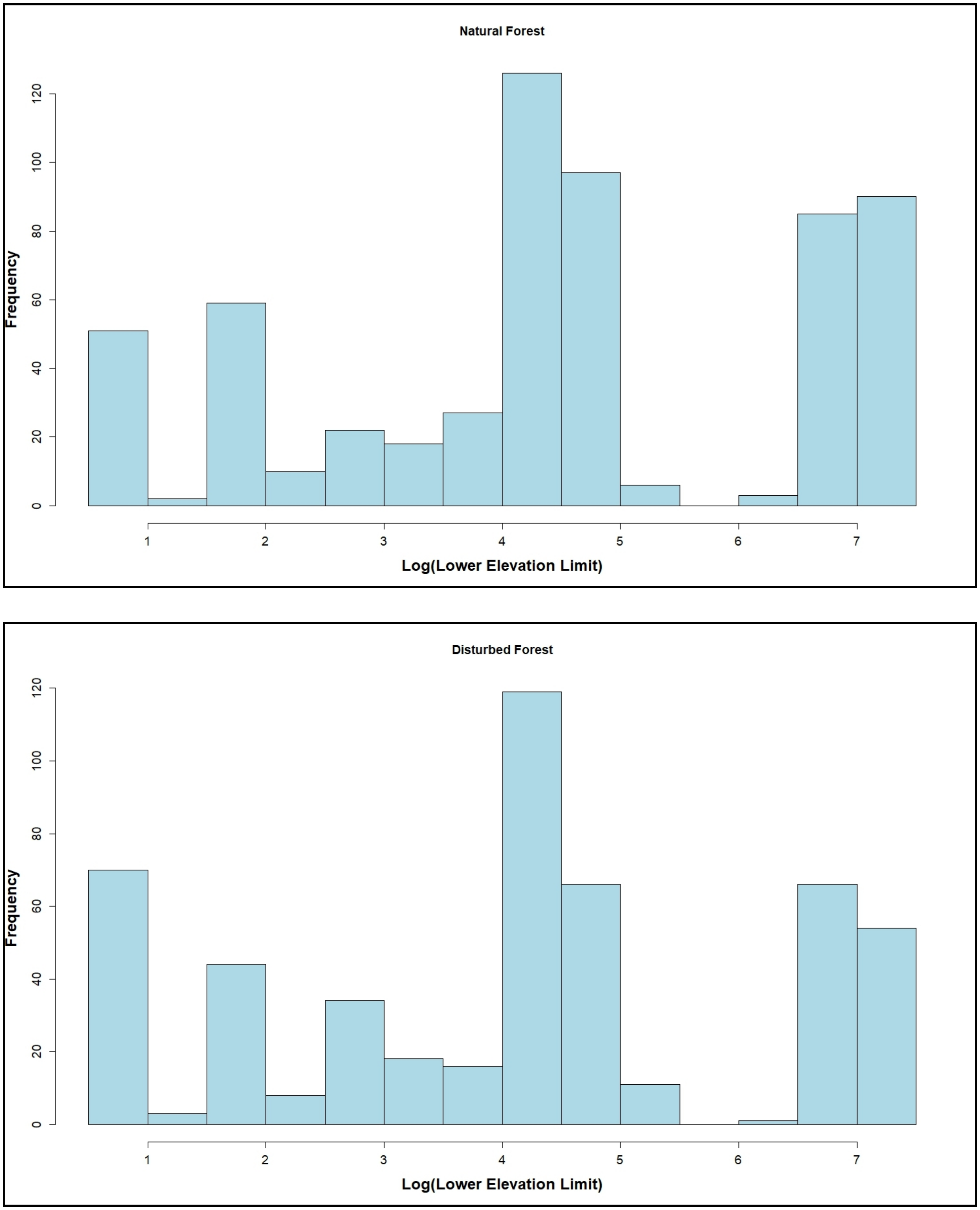


Figure S2. Site-wise distribution of host species across elevational specialisation in natural (above) and disturbed (below) forests.

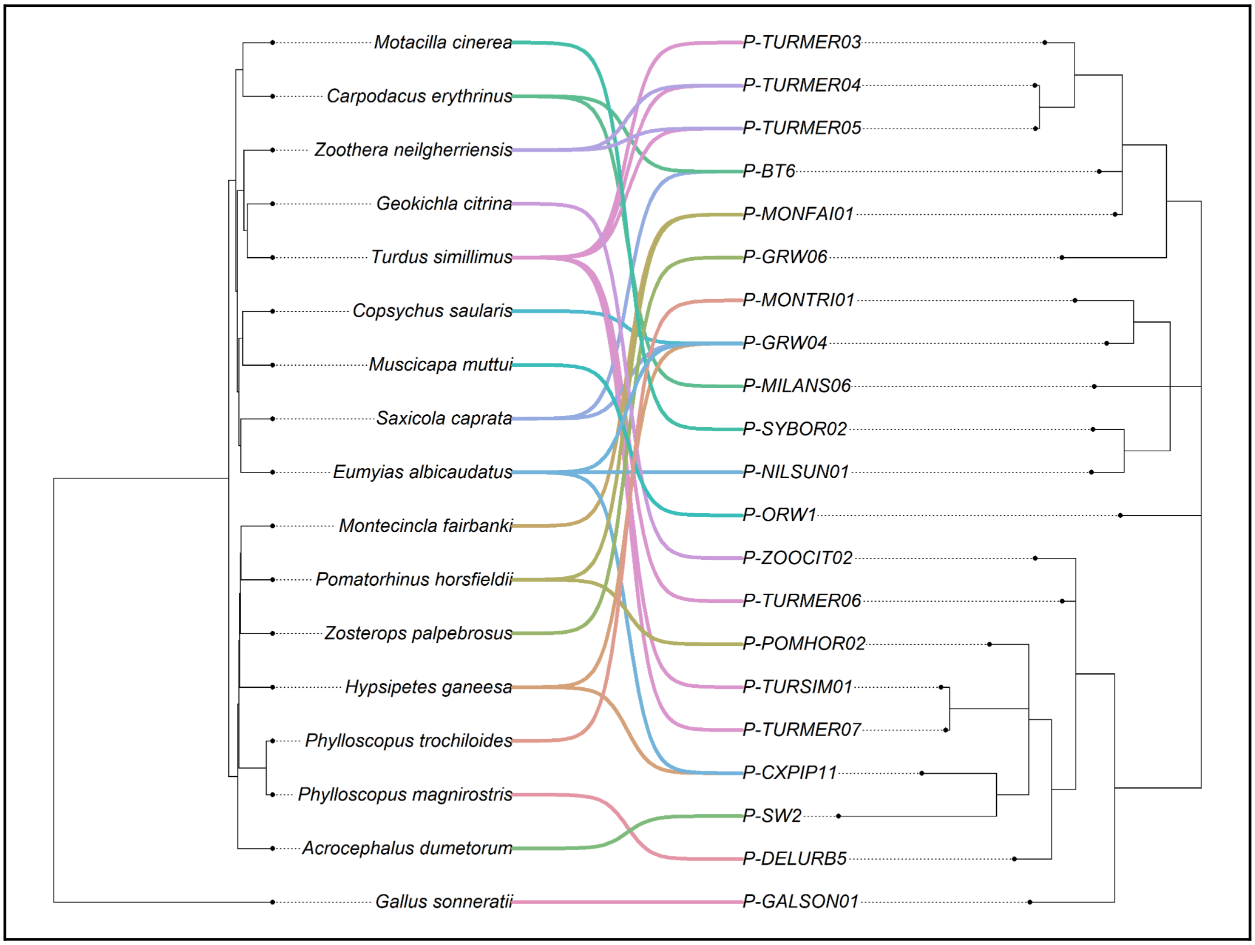


Figure S3: Host-parasite co-phylogeny of *Plasmodium* lineages.

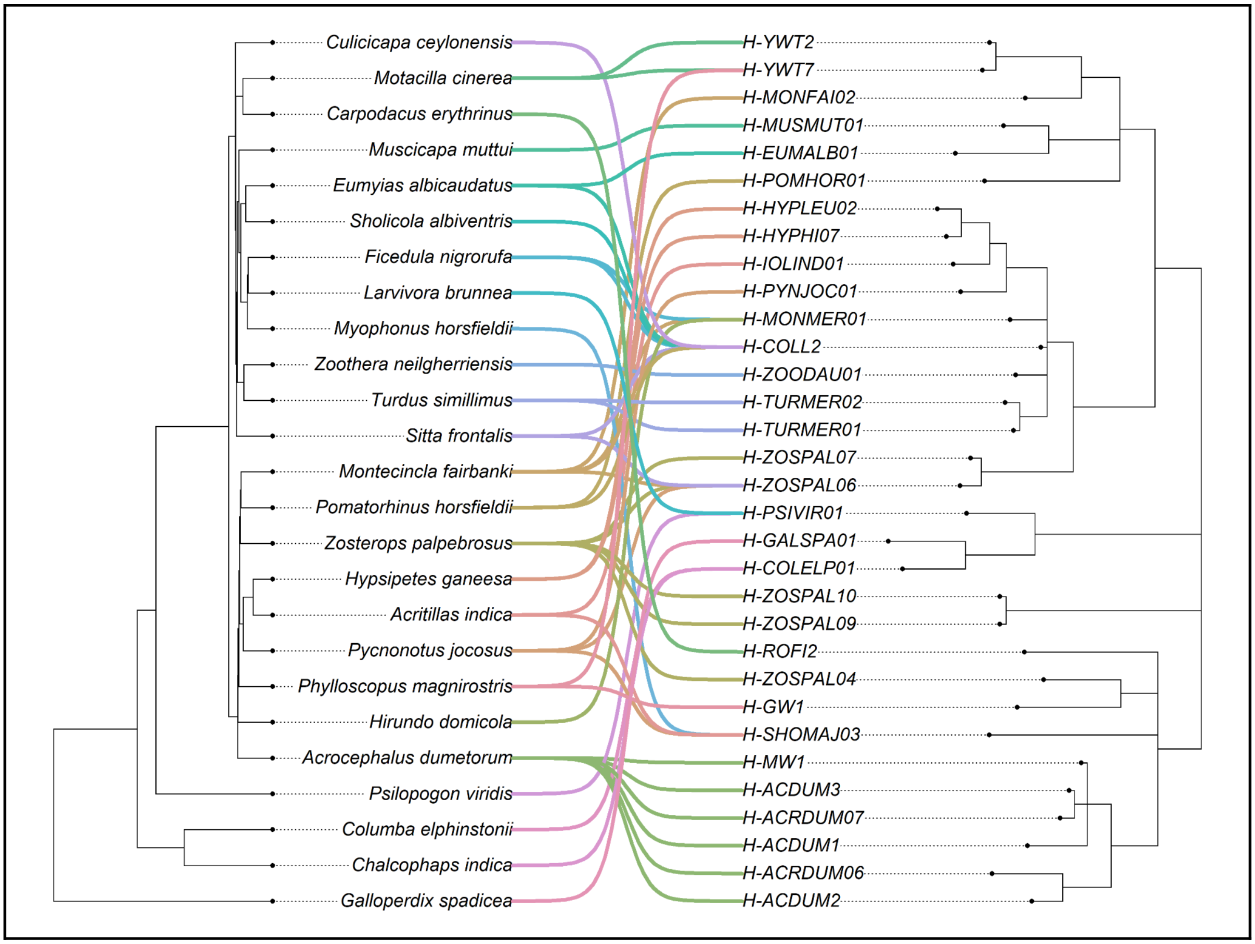


Figure S4: Host-parasite co-phylogeny of *Haemoproteus* lineages.
